# ADAR2 inhibits ER-targeted mRNA translation via regulating nucleolar assembly of signal recognition particle

**DOI:** 10.1101/2023.08.01.548241

**Authors:** Rui Feng, Yuci Wang, Dongmei Ran, Ying Li, Hui Zhou, Qi Liu, Yingchun Zhang, Honghui Ma, Hao Chen

## Abstract

Adenosine-to-inosine RNA editing is mediated by ADAR family proteins and plays critical roles in regulating gene expression. Unlike other ADAR family members, ADAR2 mostly resides in nucleolus. The exact roles of this peculiar cellular distribution of ADAR2 is still largely unknown. Here, our current study unexpectedly uncovered that nucleoli-localized ADAR2 is involved in negatively regulating ER-targeted mRNA translation, which is mediated by SRP. Importantly, this regulation is not archived through direct binding of mRNA by ADAR2 or dependent on A-to-I editing of 7SL RNA. Further analysis indicated that ADAR2 directly interacts with SRP protein components, especially SRP68, in a RNA-independent manner and probably represses the SRP assembly. Hence, we proved insights into the novel functions of ADAR2 and long-lasting enigma about the peculiar translocation of SRP proteins into nucleoli during SRP biogenesis.

## Main Text

Since the ADARs (adenosine deaminases that act on RNA) were identified around 1991, three structurally related members of the ADAR family have been characterized in human: ADAR1 and 2, which are ubiquitously expressed in many tissues with strong expression in the neuron system; and ADAR3, which is only expressed in neuronal tissues and has been proven to be enzymatically inactive [Ref 1-–2]. Intriguingly, previous studies demonstrated that ADAR2 mostly resides in nucleolus (**Fig. 1a**) [Ref 3], which primarily serves as the site for ribosome-subunit biogenesis, including rRNA transcription, processing, modification and ribosome assembly [Ref 4]. More interestingly, it was reported that enhanced translocation of endogenous ADAR2 from the nucleolus to the nucleoplasm increases the editing activity of ADAR2 [Ref 5]. However, little is known about whether there is additional biological relevance of the peculiar spatial distribution of ADAR2, especially with respect to ribosome biogenesis and protein synthesis.

**Fig. 1.**
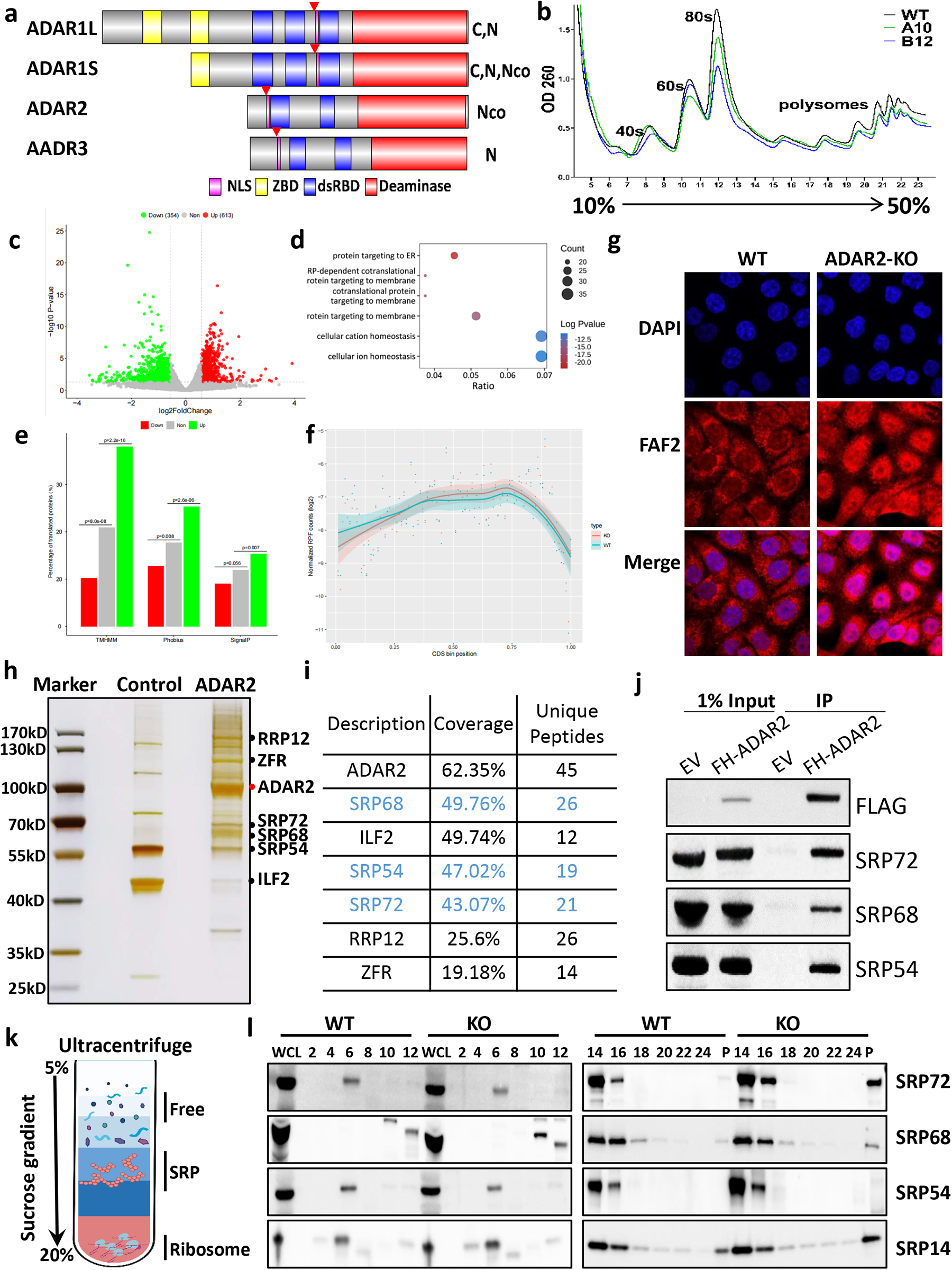
ADAR2 inhibits ER-targeted mRNA translation via direct interaction with protein components of SRP particles. **a)** A schematic domain structures of ADAR family proteins. The green box indicates the ZBD domain.The blue box indicates dsRBD domain and the red box depicts putative deaminase domain. Conserved nuclear localization signal (NLS) was shown in a purple box and highlighted with red star symbol. **b**) Polysome profiling of WT and ADAR2 KO HCT116 cells; **c**) Ribo-seq identified the translationally changed genes upon ADAR2 KO; **d**) GO analysis showed that downregulated genes is mainly enriched in ER-related functions; **e**) Phobius, SignalP, TMHMM analysis of downregulated genes-encoded proteins; **f**) accumulation of RPFs at the beginning of the respective CDS; **g**) IF showed that the protein level of membrane-localized FAF2 is significantly decreased in ADAR2 KO cells; **h**) Silver staining of ADAR2 complex purification; i) List of LC-MS/MS identified components of ADAR2 complex; **j**) the distribution of different SRP components in sucrose gradients; **k**) western blot analysis of SRP proteins in different fractions.

To uncover the potential involvement of ADAR2 in mRNA translation, we first employed the CRISPR technology to generate the HCT116 ADAR2-knockout cell lines and monitored the translatomic alteration through polysome profiling and Ribo-seq (Ribosome profiling) (**Supplementary information, Fig. S1a-c**). As the ADAR2 depletion will only affect the RNA levels of very few genes (40 upregulated genes and 59 downregulated genes), we determined to focus on investigating the translational alteration caused by loss of ADAR2 (**Supplementary information, Fig. S2a**). Considering with its nucleolar localization, we determined to employ the polysome profiling to reveal the influence of ADAR2 depletion towards global translation, which showed a very minor change upon ADAR2 knockout (**Fig. 1b**). Next, we set out to identify the transcripts that are specifically affected by loss of ADAR2 through Ribo-seq. Unexpectedly, it turns out that ADAR2 knockout significantly influences the cellular mRNA translation. A total of 1367 significantly altered events (translational efficiency: 600 up-regulated vs 767 down-regulated) were identified after comparing the translatome of WT and ADAR2 KO cells (**Fig. 1c, Supplementary information, Fig. S2b-c**). To define the biological processes, cellular locations and molecular functions that are impacted by ADAR2, we carried out gene ontology (GO) analysis of affected transcripts. Interestingly, upregulated mRNAs are enriched in GO terms of transcripts encoding membrane-localized and ER-localized proteins (**Fig. 1d**), which are dependent on the signal reorganization particle for protein synthesis [Ref 6]. By contrast, downregulated transcripts are not related with the any cellular components (**Supplementary information, Fig. S2d**). Consistently, more signal peptide-containing proteins were identified among the translationally upregulated transcripts using three different tools (including Phobius, SignalP, TMHMM) (**Fig. 1e**). Furthermore, a specific accumulation of RPFs at the beginning of the respective CDS (**Fig. 1f**), consistent with the observation that signal peptides are typically contained at the termini of respective proteins [Ref 7-–8]. Importantly, the identified altered events could be readily validated by immunofluorescence staining of a representative protein FAF2, which showed that the protein synthesis is remarkably increased in ADAR2 KO cells compared with wildtype ones (**Fig. 1g, Supplementary information, Fig. S2e**). Taken together, the above results imply that ADAR2 is likely involved in regulating the SRP-mediated translation.

We then asked about the underlying mechanism of ADAR2 regulating the functions of SRP. Considering the importance of A-to-I editing mediated by ADAR2 in RNA fates decision, we hypothesized that ADAR2 regulates these mRNA through direct interaction and RNA editing. The PAR-CLIP (Photoactivatable Ribonucleoside-Enhanced Crosslinking and Immunoprecipitation) sequencing identified the transcripts targeted by ADAR2. Consistent with previous reports, ADAR2 mainly binds with introns of pri-mRNA regions and the 3’ UTR of mRNAs (**Supplementary information, Fig. S3a & b**). However, we didn’t observe any noticeable TE changes between ADAR2-bound and unbound mRNAs (**Supplementary information, Fig. S3c**), which indicated that ADAR2 unlikely controls translation of these mRNA via direct interaction. SRP is an evolutionarily conserved ribonucleoprotein complex comprising of five SRP proteins and the RNA polymerase III-encoded 7SL RNA (**Supplementary information, Fig. 4a**), which enters nucleoli and is processed before maturation. More interestingly, mammalian Alu sequences were found to be derived from 7SL RNA in evolution and are processed 7SL RNA genes [Ref 9]. Therefore, we wondered that 7SL RNA is a novel substrate of ADAR2. To answer this question, two putative A-to-I editing site(s) were identified by analyzing transcriptomic data with very low ratio (around 0.6% – 0.9%) (**Supplementary information, Fig. 4b-d**) [Ref 10]. Next, we analyzed the A-to-I editing site(s) existing in 7SL RNA in our cell model. However, we were unable to detect any significantly altered editing events in 7SL RNA between WT and KO cells (**Supplementary information, Fig. 4e**), which implied that it’s unlikely that the ADAR2 regulates SRP function via direct RNA editing.

To uncover the underlying mechanism of ADAR2 regulating ER-targeted mRNA translation, we sought to characterize the proteins interacting with ADAR2 through ADAR2-complex purification in established stable cell line (**Supplementary information, Fig. 5a**). Interestingly, abundant protein components of SRP particles were identified in ADAR2 complex, besides the known proteins (e.g., ILF2, RPL12 and ZFR) bound with ADAR2 (**Fig. 1h-i**). Consistent with the MS/MS results, the interaction was readily validated with specific antibodies against SRP72, SRP68 and SRP54 (**Fig. 1j**). More importantly, the interaction between ADAR2 and SRP68 is RNA-independent manner (**Supplementary information, Fig. 5b**). Furthermore, we found that the N-terminus but not C-terminus of ADAR2 mediates the interaction between ADAR2 and SRP72 (**Supplementary information, Fig. 5c**). Finally, we decided to define the effects of interaction between ADAR2 and SRP proteins to the SRP functions through regulating the SRP biogenesis and found that loss of ADAR2 strikingly affects the SRP assembly and loading of SRP into polysome (**Fig. 1k-i, Supplementary information, Fig. 5d**), implying that ADAR2 is involved in negatively controlling the SRP functions.

In summary, our current study unexpectedly uncovered that nucleoli-localized ADAR2 is involved in negatively regulating ER-targeted mRNA translation, which is mediated by SRP. Importantly, this regulation is not archived through direct binding of mRNA by ADAR2 or dependent on A-to-I editing of 7SL RNA. Further analysis indicated that ADAR2 directly interacts with SRP protein components, especially SRP68, in a RNA-independent manner and probably represses the SRP assembly **(Supplementary information, Fig. 6**). Hence, we proved insights into the novel functions of ADAR2 and long-lasting enigma about the peculiar translocation of SRP proteins into nucleoli during SRP biogenesis.

## Acknowledgements

We would like to thank all members in Hao Chen’s Lab for their help and advice in experimental design. The authors would also like to acknowledge the technical support from Hua Li and Lin Lin at SUSTech CRFT. This work was supported by Center for Computational Science and Engineering at Southern University of Science and Technology. This work was supported by National Key Research and Development Program of China (2022YFC2702705), National Natural Science Foundation of China (32170604), Pearl River Recruitment Program of Talents (2021QN02Y122) and Department of Health of Guangdong Province (B2021032) to H.C. This work was also supported by Shenzhen Key Laboratory of Gene Regulation and Systems Biology (Grant No. ZDSYS20200811144002008) from Shenzhen Innovation Committee of Science and Technology.

## Author contributions

H.C., H.H.M. and Y.C.Z. conceived and designed the study. R.F., Y.L. and H.Z. performed most of the experiments. Y.C.W. and D.M.R. performed the bioinformatics analysis under the supervision of Q.L. H.C. wrote the manuscript with inputs from all other authors. All authors read and approved the final manuscript.

## Supplemental Figure legend

**Fig. S1.**
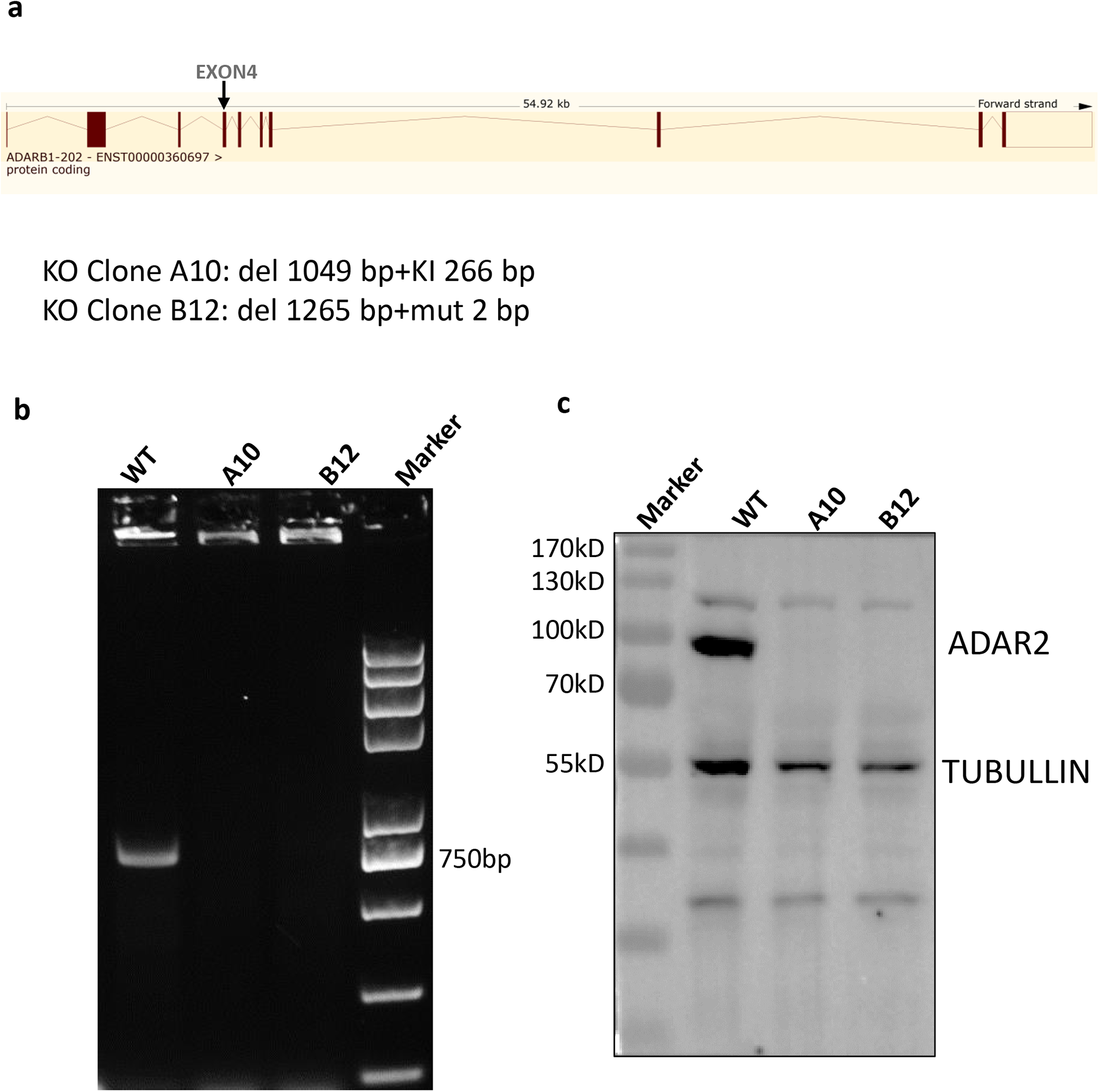
Generation of ADAR2 KO cell lines. **a)** design of ADAR2 KO in human colon cancer cell line; **b-c)** validation by PCR (**b**) and western blot (**c**);

**Fig. S2.**
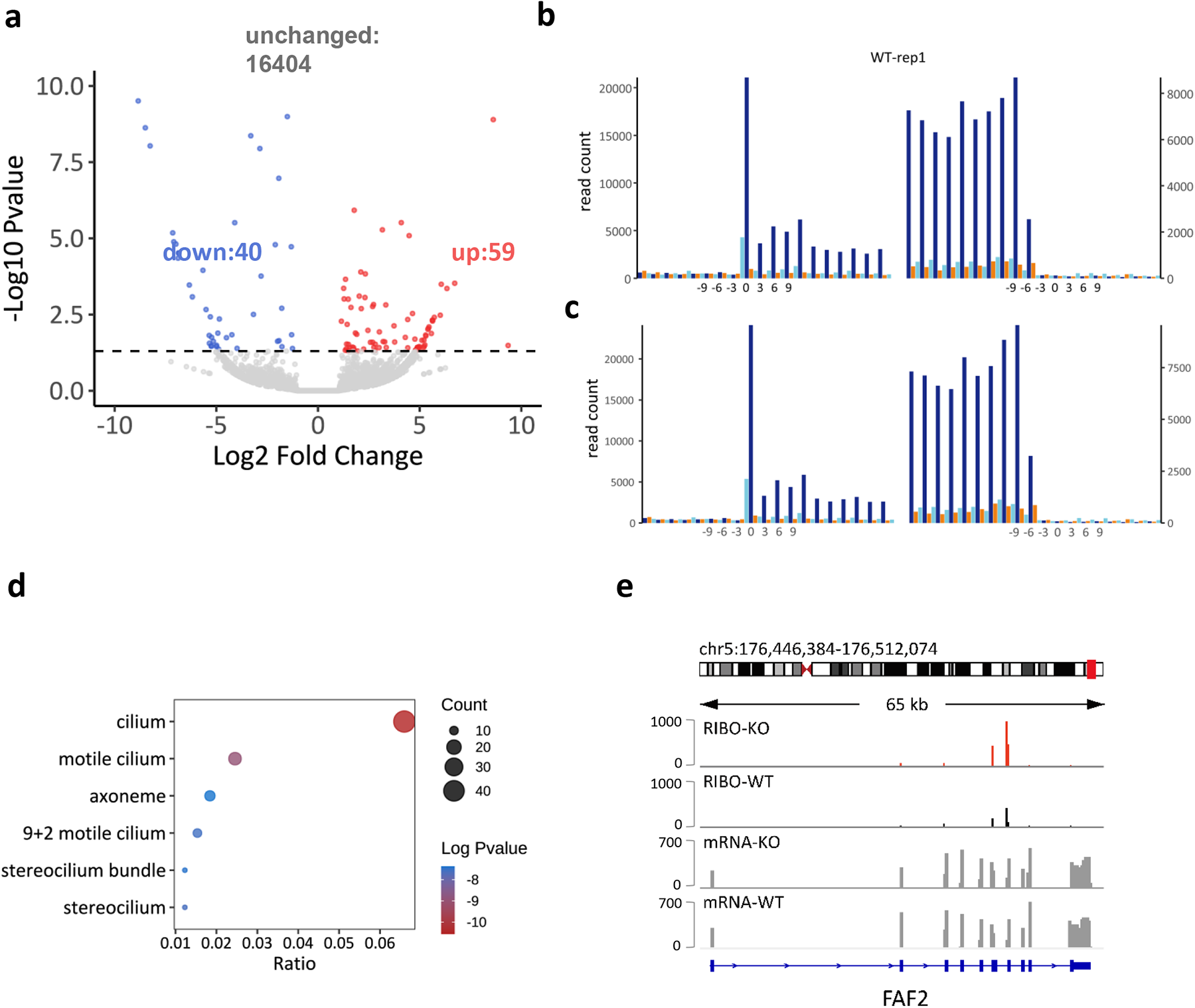
Analysis of individual mRNA translation changes by Ribo-seq. **a)** Transcriptomic analysis of gene expression; **b-c**) Quality control of Ribo-seq in WT and KO cells; **d**) GO analysis of translationally upregulated genes; **e**) Snapshot of FAF2 gene expression changes uncovered by Ribo-seq and RNA-seq;

**Fig. S3.**
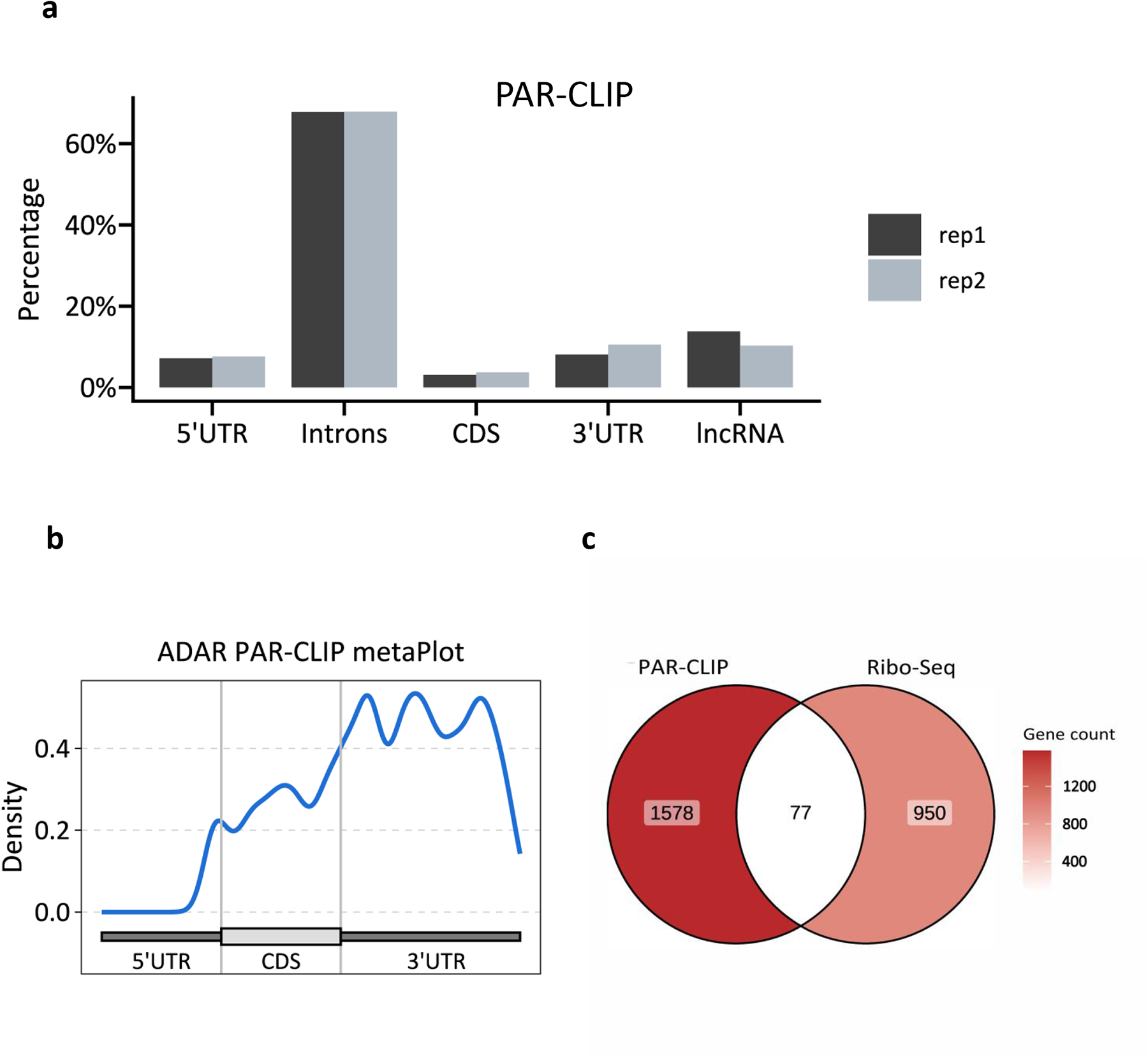
PAR-CLIP analysis of ADAR2-bound transcripts. **a**) Transcriptome-wide characterization of ADAR2 binding RNA showed that ADAR2 mainly binds with introns; **b**) the binding sites of ADAR2 are predominantly localized on 3’ UTR of mRNA; **c**) the translationally changed transcripts are seldomly overlapped with ADAR-bound ones;

**Fig. S4.**
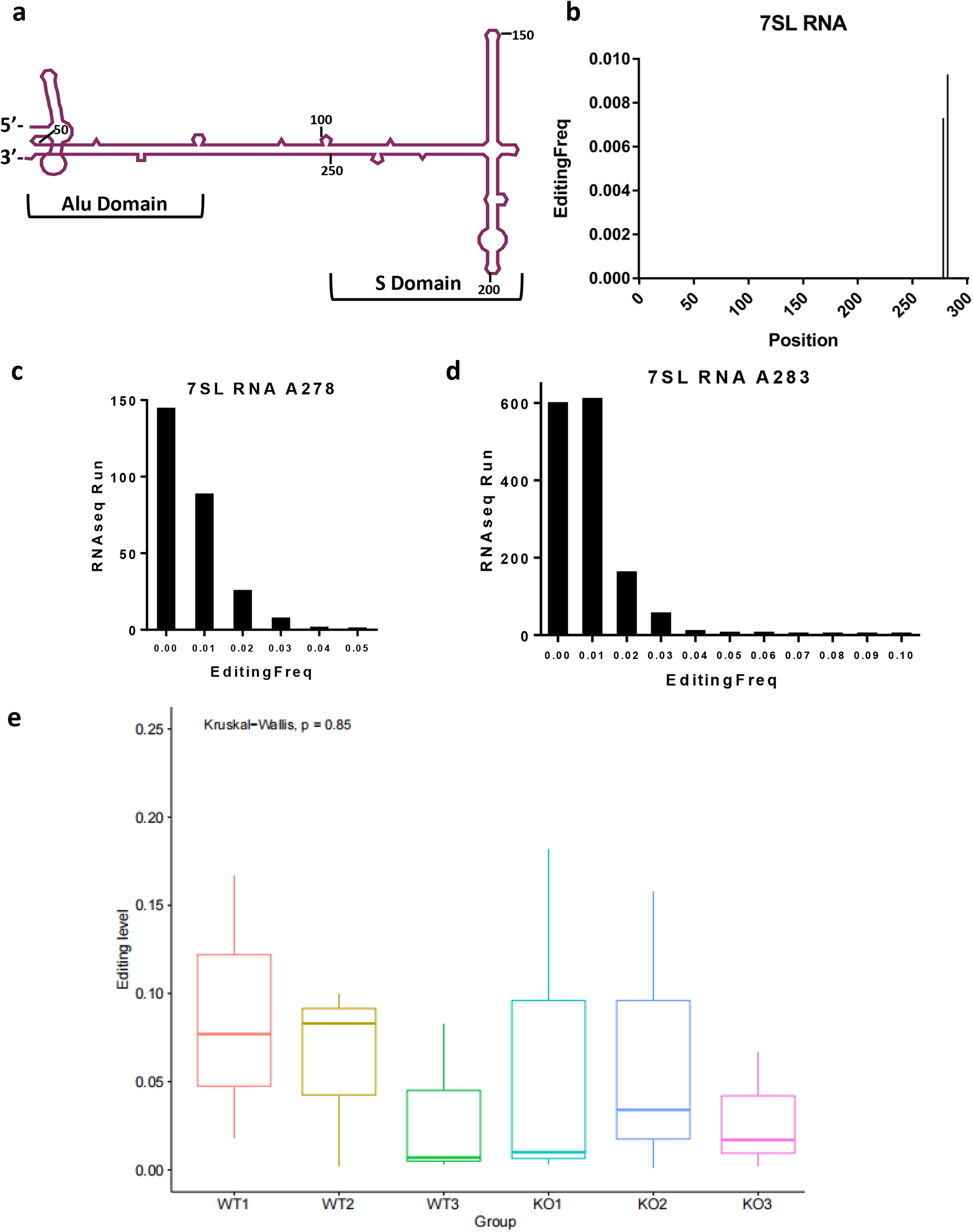
ADAR2 doesn’t directly edit SRP RNA component – 7SL RNA.

**Fig. S5.**
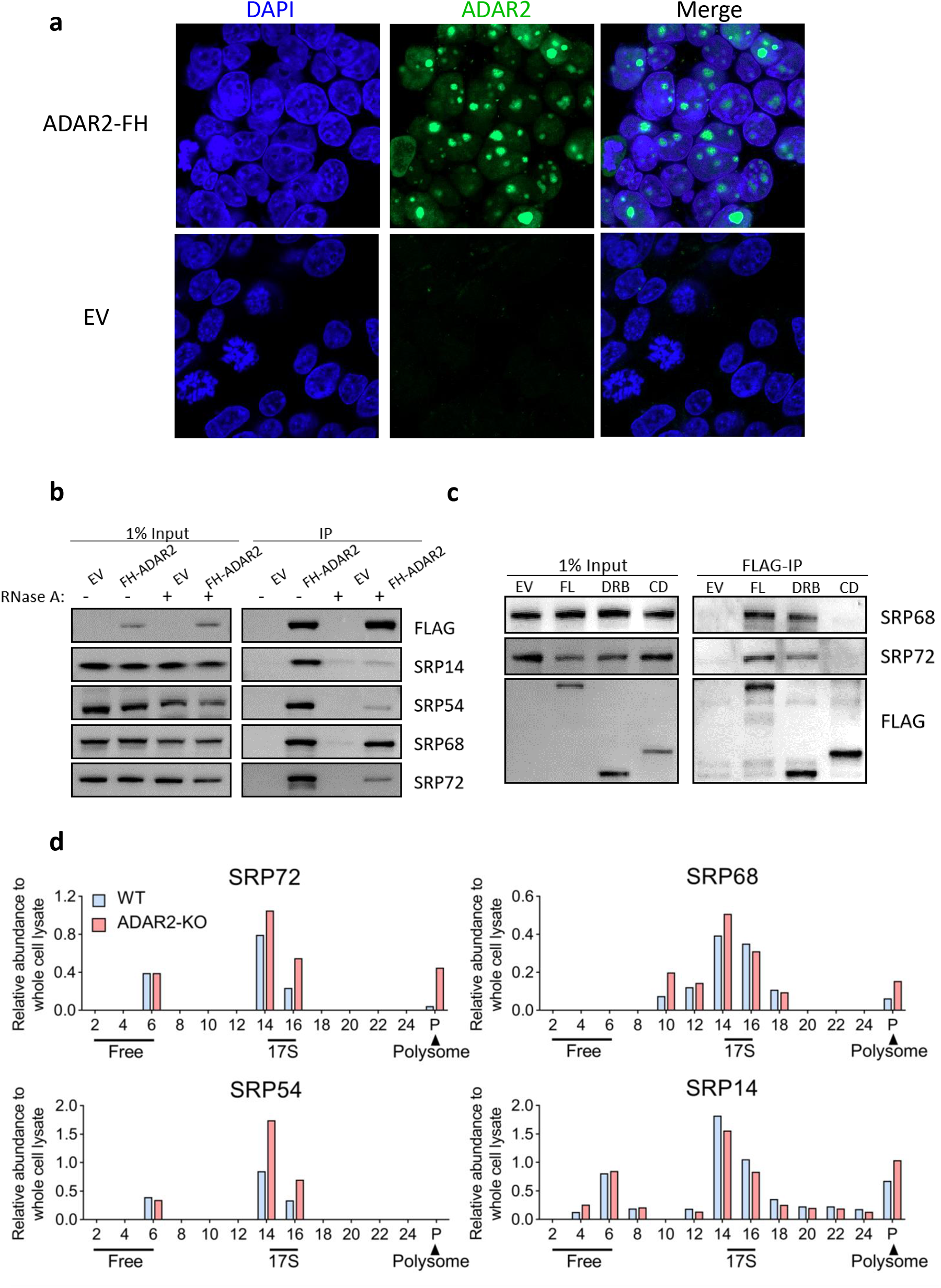
ADAR2 interacts with SRP protein components. **a**) Establishment of ADAR2 stable cell lines; **b**) ADAR2 binds with SRP 68 is not dependent on RNA; **c**) N-terminus of ADAR2 mediates the interaction between ADAR2 and SRP68 and SRP72; **d**) Relative quantification of SRP proteins in different factions;

**Fig. S6.**
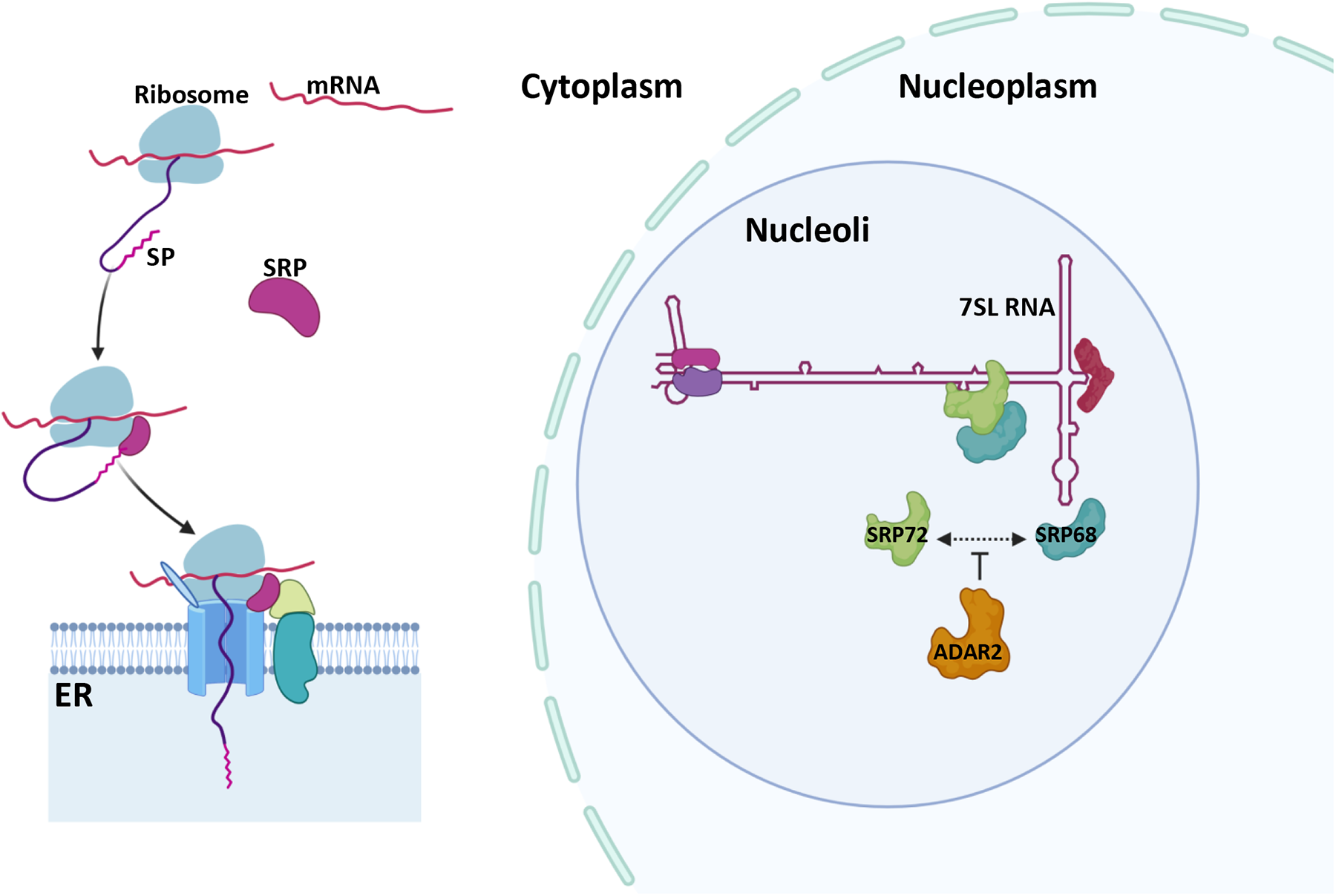
Working model for ADAR2 regulating SRP functions. ADAR2 negatively regulates ER-targeted mRNA translation. Moreover, this regulation is not archived through direct binding of mRNA by ADAR2 or dependent on A-to-I editing of 7SL RNA, while ADAR2 directly interacts with SRP68 in a RNA-independent manner and probably represses the SRP assembly.

